# Conspecific brood parasitism in the barn swallow (*Hirundo rustica*) and other Hirundinidae

**DOI:** 10.1101/2025.01.31.635828

**Authors:** Václav Jelínek, Lisandrina Mari, Adéla Petrželková, Jana Albrechtová, Jaroslav Cepák, Sylvia Kuhn, Pavel Munclinger, Oldřich Tomášek, Michal Šulc, Tomáš Albrecht

## Abstract

Conspecific brood parasitism (CBP) has been reported in many altricial avian species, yet its prevalence and underlying behavioural mechanisms remain poorly understood. We studied CBP in the barn swallow (*Hirundo rustica*), a species in which conspecific brood parasitism has been reported. We conducted parentage analysis on 1945 barn swallow broods involving 7816 offspring. Samples were collected over 12 breeding seasons at 5 colonies/populations. Contrary to expectations, we identified only six cases of CBP (0.3 % of broods). By identifying all individuals involved, i.e. the parasitic females, the female hosts and the males that fathered the parasitic eggs, we determined these cases of CBP resulted most likely from either a failed nest take-over (three times), the disappearance of a female partner or a disruption caused by researchers while only one case could be interpreted as a result of females’ mixed reproductive tactic. Based on a review of the literature on CBP in seven other Hirundinidae, we conclude that the only reliable evidence for CBP comes from cliff swallows (*Petrochelidon pyrrhonata*). Studies on other species either failed to identify the parasitic females or do not present sufficient evidence supporting the occurrence of CBP. Several studies relied on the appearance of two eggs in a nest within 24 hours to conclude that CBP occurred. However, based on the parentage analysis, we show that CBP only occurred in one out of 11 such cases in our barn swallow data. Our findings highlight that CBP is rare in our barn swallow populations, and have been likely overestimated in other studies. We emphasize the importance of frequent nest checks and rigorous genetic validation in CBP research. Overall, our results challenge existing assumptions about the prevalence of CBP and provide insights into the behavioural mechanisms that lead to CBP, with ecological and evolutionary implications.

## INTRODUCTION

Given that parental care is costly (Clutton-Brock 1991; Kölliker 2012), individuals can increase their fitness by exploiting the care of others. For female birds, one way to achieve this is to lay eggs in the nest of other females, a reproductive strategy referred to as brood parasitism. Obligate brood parasites, such as cuckoos and cowbirds never build a nest or rear their own offspring, but instead rely on the care by the parents of the host species (Davies 2010). About 1% of all bird species belong to this category. In contrast, conspecific brood parasitism (CBP), in which some individuals incubate their own eggs whereas others lay eggs in the nests of other individuals of the same species, or in whic an individual lays some eggs in its own nest and some in the nest of a conspecific, has been reported in at least 2.5 % of all bird species (Yom□ Tov 2001; Yom-Tov and Geffen 2017).

CBP has evolved multiple times and is particularly prominent in precocial bird species, especially among Anatidae (Yom-Tov and Geffen 2017). In contrast, CBP is less frequent in altricial species and is mostly restricted to colonial or semi-colonial species, such as the Ploceidae, Sturnidae or Hirundinidae (Yom□ Tov 2001; Weaver and Brown 2004; Yom-Tov and Geffen 2017). This difference between precocial and altricial species is likely due to differences in opportunities to parasitize and in the costs of parasitism to the host parents associated with different life histories. Unlike precocial species, which typically have large clutches and lower parental investment per offspring, altricial species tend to have small clutches, often start incubation during egg-laying, and provide more intensive parental care to their offspring (Ar and Yom-Tov 1978; Andersson 1984; Sayler 1992; Sorenson 1992; Lyon and Everding 1996). In both precocial and altricial species, enlarged clutches may increase the risk of hatching failure (Eadie 1989; Semel and Sherman 2001) or predation (assuming it prolongs the incubation period; Gibbons 1986; Nielsen et al. 2006). In altricial species, the presence of a parasitic chick can also increase competition within the nest, further reducing the number and quality of host offspring (Brown and Brown 1991). Moreover, in both groups, host fitness may be reduced if the parasitic female damages or removes some of the host’s eggs (Lombardo et al. 1989).

CBP has been considered an alternative reproductive strategy with potential adaptive benefits for parasitic females (e.g., Sandell and Diemer 2010, Reichart et al. 2010, Monclús et al. 2017). These benefits can reflect female’s physical state or accidental events, such as nest loss. Parasitic female may lay eggs in the nest of another female after losing her own nest (to avoid having zero reproductive success in a given season), increase her overall reproductive success by laying eggs both in her own nest and in that of another female or some females may specialize in reproducing via CBP either lifelong or during a specific phase, similar to obligate brood parasites (summarized in Lyon and Eadie 2008).

Distinguishing between these scenarios requires information about the focal nest and its owners, and about the parasitic individual. Molecular parentage analysis allows researchers to identify individual females’ offspring spread across multiple nests within a population (Reichart et al. 2010, Tiedemann, et al. 2011). However, determining the occurrence of CBP requires a combination of genetic methods, observations of marked individuals, and frequent nest checks during egg-laying (given that female birds can only lay one egg per day Welty 1963; Brown 1984; Sturkie 2012). Such combined approach can answer important questions about the evolution of CBP and the circumstances under which it occurs (Griffith et al. 2004; Lyon and Eadie 2008; Brown and Brown 1988, 1989).

One species in which CBP has been reported to occur frequently is the barn swallow (*Hirundo rustica*). In a four-year study, Møller (1987) reported CBP in 43 out of 261 (16.4%) breeding attempts, based on the appearance of two eggs in the nest on the same day. Parasitic eggs were identified by comparing the eggs in the female’s own nest with those found in neighbouring nests. However, later study revealed that human ‘naked-eye’ assessment is not a reliable method for inferring brood parasitism in this species (Brown and Sherman, 1989) and in the closely related cliff swallow (*Petrochelidon pyrrhonota*; Brown and Sherman, 1989). Even in species with much larger inter-individual variation in egg size and appearance, identifying individual females based solely on egg characteristics is difficult (McRae 1997, Grønstøl et al. 2006, Šulc et al. 2022). These findings highlight the need for follow-up studies using parentage analysis to identify the prevalence of CBP (Hughes et al. 2024).

Here, we combine nest monitoring in a colour-banded population with parentage analysis to study CBP in the barn swallow. Specifically, we used a dataset of 1945 intensively monitored breeding attempts and genotypes of 7816 nestlings and 2198 adults. This allowed us to (1) assess the frequency of CBP in our barn swallow populations, (2) identify the parasitic females, and (3) elucidate the circumstances leading to CBP. An additional aim of our study was to test the relationship between colony size and the frequency of CBP. However, CBP turned out to be so rare that no meaningful test could be conducted. We also carefully and critically reviewed all studies analysing the genetic parentage in the barn swallow and all possible cases of CBP in other species of Hirundinidae species to evaluate the importance of this alternative reproductive strategy for their reproduction.

## METHODS

The barn swallow is a socially monogamous passerine species, primarily known for synanthropic breeding in large colonies in barns and stables. However, smaller colonies consisting of just a few pairs and solitary breeding are also common. We collected data from five local populations of varying sizes and spatial structures in Southern and Eastern Bohemia (Czech Republic) between 2010 and 2021 (Table 1).

**Table 1:** Characteristics of the studied barn swallow populations in Southern and Eastern Bohemia, Czech Republic. For each colony/population, we give the location (coordinates), the breeding seasons during which they were studied, the mean number of breeding attempts per year and the range. Note that the Chotovice population consists of 23 individual sites with solitary pairs or small colonies with few pairs (16 sites in 2017, 19 in 2018 and 19 in 2019). We thus give the mean size per site, as well as the total number of breeding attempts at all the sites combined (between brackets).

| Population | Coordinates | Breeding seasons | Colony size |  |
| --- | --- | --- | --- | --- |
|  |  |  | Mean | Range |
| Hamr | 49.06 N, 14.77 E | 2010 - 21 | 62.4 | 43 - 102 |
| Šaloun | 49.07 N, 14.71 E | 2010 - 21 | 69.5 | 40 - 88 |
| Břilice | 49.02 N, 14.74 E | 2016 - 21 | 65.7 | 38 - 86 |
| Stará Hlína | 49.04 N, 14.82 E | 2019 - 21 | 39.3 | 31 - 47 |
| Chotovice | 49.85 N, 16.17 E | 2017 - 19 | 3.1 (55.0) | 1 - 15 (43 - 65) |

Each population was surveyed throughout the entire breeding season, from late April to early September, with varying levels of intensity depending on the year and the stage of the breeding season. Generally, visits were less frequent from early July when usually more than half of second clutches had already been laid. In seasons with minimal research activity, the interval between visits usually extended from every three to four days early in the season (May and June) to seven or more days toward its end (July and August). During periods of maximum research intensity, daily visits were conducted until mid-July, followed by a gradual reduction in effort thereafter. Chotovice population was surveyed even less intensely to maintain good relationships with homeowners. One-pair localities were visited three or four times per season while localities with more pairs were visited more frequently. During each visit, nests were carefully checked and their content recorded, while newly build nests were actively searched for. Newly laid eggs were marked using felt tip pen during six breeding seasons (2013–2015 and 2018–2021) to collect data on precise laying sequence.

During each breeding season at each colony, we tried to locate all nests and collected a blood or tissue sample from all nestlings and from all unhatched eggs. We sampled 7816 offspring (nestlings or eggs) from 1945 nesting attempts while 315 nesting attempts with at least 1075 offspring were not sampled mainly due to complete predation or nest destruction. On sampled nests, 982 offspring were not sampled. These offspring consisted of infertile eggs, eggs destroyed during handling, and eggs or nestlings that disappeared from their nests due to death, partial predation, infanticide, or other causes. At each location, we also captured adult birds throughout the season using mist nets or custom-made landing nets. We sexed each adult in the hand based on the shape of their cloacal protuberance. We also marked each adult with a unique combination of a standard aluminium ring and up to three coloured plastic rings, allowing us to identify nest owners using binoculars or cameras.

We took a blood sample (∼20 μl) through venipuncture from all live adults (N = 2186) and nestlings (N = 7190), and a tissue sample from embryo material from unhatched eggs (N = 468) and from dead nestlings (N = 158) or adults (N = 12). All samples were stored in 96 % ethanol. DNA was extracted using various kits, primarily the NucleoSpin Blood kit (Macherey-Nagel, Germany) following the manufacturer’s instructions. We obtained a final DNA concentration of at least 20 ng/μl, which we considered the minimum for accurate allele determination. Low-yield samples, such as those from unhatched embryos, were concentrated using magnetic beads (Ampure XP beads, Beckman Coulter) to increase their DNA concentration. If the DNA concentration remained below 20 ng/μl after the concentration step, the number of PCR amplification cycles was increased by up to six cycles (see below). A total of 2198 individual adults (1207 males and 991 females) were included as candidate parents across all analyses.

Individuals were genotyped at 15 highly-polymorphic autosomal microsatellite markers and at 2 sex-specific markers, which were amplified in two multiplex reactions (Table S1). One sex-specific locus was included in each multiplex mix to confirm the sex of each individual and as an internal control. 329 out of 765 samples from the Chotovice area were genotyped at a total of 15 loci (including one sex-specific locus). Primers were fluorescently labelled (see Table S1), and PCR reactions were carried out using the Qiagen Type-It Microsatellite PCR kit. PCRs were conducted in a total reaction volume of 10 µl, containing 1 µl of DNA (20-80 ng/µl), 1 µl of one of the two primer mixes, 5 µl of Qiagen Type-it Microsatellite PCR Master Mix, and 3 µl of PCR-grade water. The PCR conditions were as follows: initial denaturation for five minutes at 95°C, followed by 30 cycles of 30 seconds at 94°C, 90 seconds at 50°C (for Mix 1) or 52°C (for Mix 2), 60 seconds at 72°C, and a final extension step of 45 minutes at 60°C. We added 1.5 µl of the PCR product to 13 µl of Hi-Di formamide and GeneScan™ 500 LIZ™ Dye Size Standard (ThermoFisher Scientific), denatured the mixture for five minutes at 95°C, and loaded it onto an Applied Biosystems 3100 or 3130xl BioAnalyzer (ThermoFisher Scientific).

Genotypes were determined using Geneious Prime® 2024.0.3 software (GraphPad Software LLC d.b.a Geneious), with allele determination (peak detection) performed blindly by two different persons (VJ, LM and three technicians) to ensure accuracy. To create allele bins, all fragment lengths for a given marker were plotted in MS Excel, from shortest to longest. Gaps between neighbouring alleles were then identified, and final bins for each allele were manually created in Geneious, using the shortest and longest PCR fragment of one set that were assumed to have amplified from the same allele as the border values for each bin. Finally, all individuals were genotyped in Geneious with the binning information.

Parentage assignments for all offspring samples were conducted using Cervus version 3.0.7 (Field Genetics Ltd.), following the methods outlined by Kalinowski et al. (2007). Given that the sex of each adult was known with certainty, we carried out parent-pair analysis. Adults were genotyped at all 17 loci, and offspring samples were included in the analysis if they had been genotyped at a minimum of 10 loci (98.7% of offspring samples were genotyped at all loci). When the same sample or individual was genotyped multiple times, their genotypes were compared, and any discrepancies were corrected. The combined non-exclusion probability for the first parent was > 3.93 × 10^−6^ and the non-exclusion probability for the second was > 3.71 × 10^−9^ in all 15 parentage analyses performed to assign maternity.

Parentage analysis was conducted in four sequential steps for each breeding season. Step 1: we included only candidate parents captured or observed during a given breeding season. Step 2: we included adults from both the previous and subsequent seasons to identify parents for offspring which were not assigned in the first step. Step 3: we included adults observed two years prior and two years after the target season as candidate parents. We used this stepwise approach instead of a single large analysis including all candidate parents across five seasons to minimize the risk of false assignments, which could arise due to the presence of related individuals in our populations. Notably, despite high recruitment rates (with up to 25% of the male population consisting of nestlings from the previous year), no correctly assigned parent from one step was ever replaced by another in a subsequent step.

Parentage analysis was conducted in four sequential steps for each breeding season. Step 1: we included only candidate parents captured or observed during a given breeding season. Step 2: we included adults from both the previous and subsequent seasons to find parents for offspring which was not assigned in the first step. Step 3: we included adults observed two years prior and two years after the target season as candidate parents. Step 4: we used paternity analysis with known mothers as a control for situations when Cervus incorrectly assigned mothers, when the fathers’ genotypes was unknown. This used to happen even though the genotype of correct but unassigned mother perfectly matched the genotype of the offspring (N = 7, genotypes were checked manually). As known mothers we chose mothers which were correctly assigned (see below) by the Cervus in step 3 for all or the majority of offspring in each nest. We used this stepwise approach instead of a single large analysis including all candidate parents across five seasons to minimize the risk of false assignments, which could arise due to the presence of related individuals in our populations. Notably, despite high recruitment rates (with up to 25% of the male population consisting of nestlings from the previous year), no correctly assigned parent (see below) from one step was ever replaced by another in a subsequent step.

A parent pair was considered correctly assigned under two circumstances. (1) If there were zero mismatches between the parental alleles and the offspring alleles. (2) If one or two mismatches occurred between putative parents and offspring, the specific allele(s) were carefully checked. If an error in allele scoring was identified, it was corrected, and the entire process of parentage analysis was repeated with the updated genotype. If no scoring error could be found, we considered whether the mismatch could be explained with the existence of a null allele. Instances of potential null alleles were documented, and parents were considered as correctly assigned. In the remaining cases of a single mismatch, the mismatch was attributed to a mutation in 1.2% of all offspring, with 53% of these mutations occurring at the Hir20 locus). In all instances when candidate for CBP was identified, we manually matched genotypes of genetic (parasitic) and host (parasitized) mothers to confirm the correct assignment.

Genetically assigned parents were then cross-referenced with the field data. We selected all cases where multiple genetic mothers were identified within the same clutch or brood as candidates for CBP. To verify the parasitic status of each candidate female, we reviewed all available information, including laying sequences, field notes, and photographs. An offspring sample was classified as the result of CBP only if we were unable to disprove its parasitic status due to any possible error or mistake during fieldwork, sampling or handling of samples and/or if the additional evidence supported the conclusion of brood parasitism.

## RESULTS

We identified six cases of putative CBP. Thus, 0.3% of broods (N = 1945) contained one or two parasitic eggs, and 0.09% of the genotyped offspring (N = 7816) were parasitic. Below, we provide a detailed description of each case.

### Cases of CBP - based on parentage analysis

#### Case 1: Šaloun 2013

In 2013, we found the first egg in nest 32 (a second breeding attempt in that nest that year) on 15 July. Daily nest checks thereafter showed that no other eggs were laid in the next three days. We found the second egg on 19 July, followed by three more eggs over the next three days, resulting in a total clutch of five eggs. All eggs later hatched. Female S590599, who was the social female, was the genetic mother of four nestlings, while female S678065 was the mother of the fifth. Female S590599 had a first breeding attempt in nest 32 with male S678070 and they raised four nestlings (all within-pair young). However, this male did not sire any offspring in the second brood. The four non-parasitic nestlings had three different genetic fathers but we were not able to assign their social father. Males S540966 and S540616 each sired one nestling, while male S644662 sired the remaining two. Interestingly, male S644662 also sired the “parasitic” nestling. The parents of this nestling (male S644662 and female S678065) bred together in nest 123 (first breeding attempt, first egg laid on 3 June), and raised four nestlings (all within-pair young). Female S678065 also had a second breeding attempt in nest 85 (first egg laid on 21 July), where she raised three nestlings sired by two different males. Her social partner in this attempt was unknown.

#### Cases 2 and 3: Hamr 2018

In 2018, we checked nest 225 daily, revealing laying of one egg each day at the turn of April and May. We found the fifth egg on 2 May at 11:30, but failed to number it. We checked the nest again on the same day at 17:00 and found a sixth egg. Genetic analysis revealed that one of the two eggs laid on 2 May (of which one hatched, while the other did not) was parasitic. The genetic parents of this unhatched parasitic egg were identified as female S952410 and male S746970, who had their own nest in a different barn 40 meters away. We checked their nest (#129) for the first time on 3 May at 9:30 and it was empty. We checked it again on 4 May at 9:30 and then it contained two eggs. Over the next four days, three more eggs were laid, resulting in a total of five eggs. Genetic analysis revealed that all five eggs were from the social parents.

This pair later bred for a second time in the same nest (#129; *Case 3*). On 20 June the nest was empty. We checked it again on 23 June when we found there two eggs followed by two more eggs laid in next two days. All eggs were sampled (three nestlings and an unhatched egg). Two nestlings and the unhatched egg had the same parents (female S952410 and male S746970), while the third nestling was parasitic sired by the same male (S746970). The mother of the nestling was female S764105, who bred in the nest 354 located one meter from nest 129, with male S952158. The only nestling which hatched from five egg clutch on nest 354 most likely fledged between 22 June and 24 June. During this time its mother laid the parasitic egg in the nest 129. Female S764105 then moved to another barn where she bred with originally unpaired male S764137 in nest 16.

#### Case 4: Hamr 2019

On 20 July 2019, female S961968 and male S764054 started a second clutch in nest 129 after a successful first breeding attempt in the same nest. We checked the nest again on 24 July and found five eggs, indicating a typical pattern of one egg being laid per day. All five eggs hatched, but one nestling disappeared from the nest before sampling. Genetic analysis of the other four nestlings showed that female S961968 was the mother of two of the nestlings (both sired by extra-pair father S952049), while female S764166 was the mother of the other two with male S995231 being their father. This latter pair had successfully bred together in nest 4118 in the same barn (first breeding attempt), but then their nest was successfully taken over by another pair (male S952418 and female S764168). After the take-over of her nest, on 24 July, female S764166 was observed in a fight at nest 129 with the female owner S961968 (Fig. 1). Figure 2 shows photos of the parasitized second clutch and the non-parasitized first clutches of both the parasitic female (S764166) and the host female (S961968).

**Figure 1.**
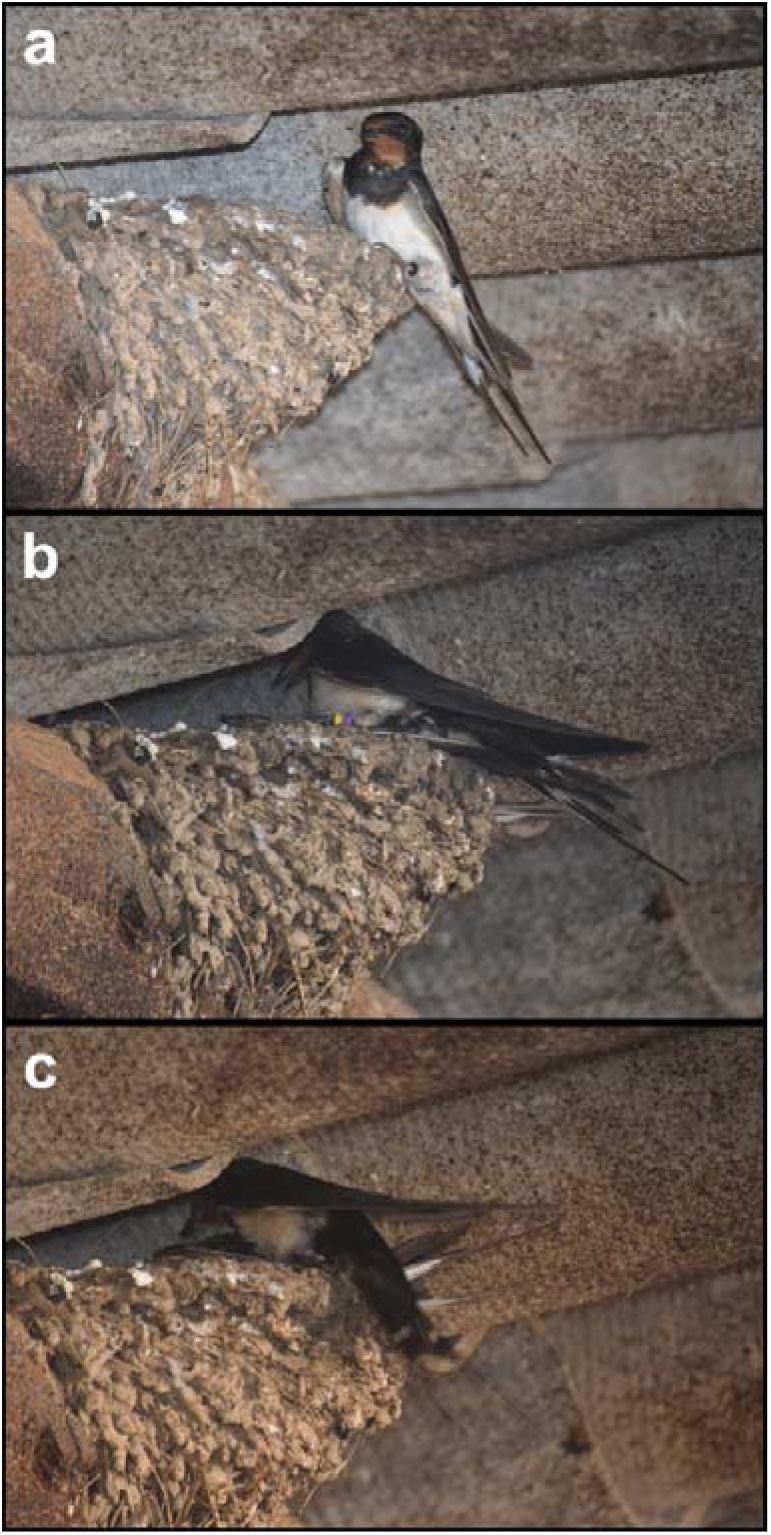
Photos showing a fight between two barn swallow females at nest 129. (a) Female S764166 (black plastic ring on the left leg) sits on the nest rim.(b) The owner of the nest, female S961968 arrives (purple above yellow ring on the left leg). (c) Both females fight.

**Figure 2.**
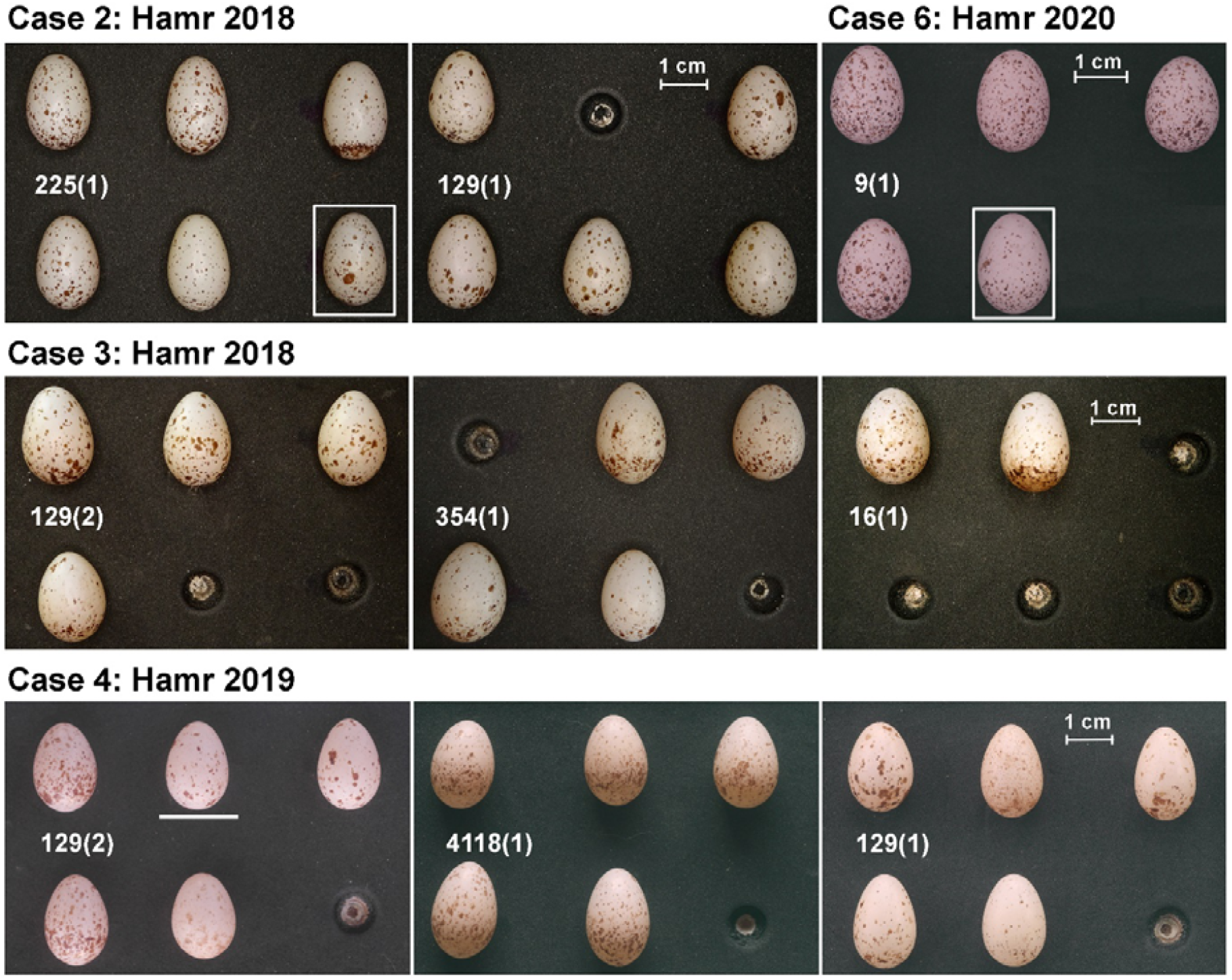
Photos of barn swallow clutches featured in cases of conspecific brood parasitism. *Case 2*: 225(1) was the first nesting attempt of host female S879473. The parasitic egg was laid by the female S952410 and did not hatch (framed). The nest 129(1) is the first nesting attempt of parasitic female S952410. *Case 6*: Parasitized nest 9(1) at Hamr colony. The fifth parasitic egg (framed) was laid and photographed later during the incubation. *Case 3*: The parasitized clutch 129(2) was the second nesting attempt of the female S879473. Three eggs belonged to this host female but one egg was laid by a parasitic female S764105 which had two more clutches in 2018, 354 (1) and 16 (1). *Case 4*: 129(2) was the second nesting attempt of host female S995231. Maternity analysis revealed that only two eggs were laid by this female. The other two eggs were laid by parasitic female S764166. The fifth egg on the photo (underscored) could not be assigned to any female because the nestling which hatched from this egg disappeared before blood sampling. The clutch 4118(1) shows the first nesting attempt of female S764166 and the 129(1) is the first clutch of female S995231.

#### Case 5: Břilice 2019

In 2019, we found the first egg in nest 6719 on 7 June and an additional egg every day until 11 June (all eggs were numbered). When we checked the nest again on 13 June, it still contained five eggs. However, during the next nest check on 24 June, we discovered a sixth, unmarked egg in the nest. On 25 June, we found that four eggs had hatched and on 26 June, we collected the remaining two eggs (numbers 4 and 6) and later dissected them. We obtained a DNA sample from egg 6 (but not from egg 4), and from two of the nestlings (all died after the parents had abandoned the nest). Genetic analysis revealed that female S877663 (nest owner) was the mother of the two nestlings, while egg 6 had been laid by female S962047. The parasitic female had her own nest (#8619), located 25 m from the focal in the same barn. She was caught in a landing net at her nest on the morning of 17 June with an egg in her belly. She was released within about 15 min of capture, after banding, measuring and blood sampling. We checked nest 8619 in the evening on the same day and found two eggs. However, we had not checked the nest in the preceding days. The final clutch size in nest 8619 was four eggs.

#### Case 6: Hamr 2020

In 2020, we found the first egg in nest #9 on 1 May and an additional egg on each consecutive day until 4 May (all four eggs were numbered). We checked the nest on 5, 6, 7 and 10 May and found four eggs (as expected), but during a following check on 17 May we discovered a fifth, unmarked egg.

Hatching started on 19 May and on 21 May we found four nestlings and the unhatched egg #5. We collected the egg and obtained a tissue sample from the embryo as the egg had only been incubated for approximately 6 or 7 days. The parasitic egg was laid by female SC26826 and sired by male SC26829. This pair had been caught together on 11 May at an empty nest (#106) located 7 m from the focal nest. The female had a brood patch (earliest stage of development). After catching, both birds disappeared from the site. The male was never recorded again, but the female was recaptured on 8 June at another colony (Stará Hlína), four kilometres from Hamr, where she stayed for two breeding seasons.

### Potential cases of CBP - nests where two eggs appeared on the same day

In 11 out of 587 breeding attempts (1.9%) for which we documented the complete egg-laying sequence, two eggs appeared within 24 hours interval. However, parentage analysis of 10 of those clutches showed evidence for CBP only in a single case (Case 2, described above). In seven cases, all eggs hatched and were genotyped. In two nests, only 1 out of 4 and 1 out of 2 eggs were genotyped (due to the disappearance of an egg and nestlings that had most likely died at an early age). In all cases, the social mother was confirmed as the genetic mother. The remaining nest was depredated before sampling.

## DISCUSSION

Genotyping of 7816 offspring from 1945 barn swallow breeding attempts revealed a low frequency of putative CBP in this species: 0.3% of broods contained an offspring that was not related to the social female at the nest (N = 6 cases). The low occurrence of CBP precluded testing the effect of colony size on CBP prevalence. We discuss the circumstances that led to the few cases of CBP and compare our results with those from other studies on barn swallow. Finally, we discuss also the evidence for CBP in all published studies of barn swallow and other Hirundinidae.

Our observations suggest that at least one case of apparent CBP occurred after a nest take-over, resulting in a contest between two pairs for occupancy of a single nest (see Results, Case 4). Nest take-over might also explain another case of CBP (Case 1), where the first egg was laid atypically three days before the second one and could have been parasitic. A possible scenario is that the parasitic female unsuccessfully tried to take over the nest, and then bred elsewhere (nest 85), with the original owner accepting the parasitic egg. Alternatively, the parasitic egg might have been laid mistakenly in this nest (which was 2 m from nest 85), as accidental egg-laying in the wrong nest does occur (Turbek et al. 2019). Each breeding season, we observe several instances in which a single egg appears in an empty nest, and is subsequently deserted. In two instances in which the egg could be genotyped, the female laying these eggs bred in a nearby nest (within 1 m), and the egg was missing from the female’s laying sequence. However, this was not the case here, as the parasitic female’s next breeding attempt (nest 85) likely started several days after the suspected parasitic egg had been laid. Similar scenario likely occurred on nest 129 (Case 3), where the parasitic egg was laid by a female who likely lost her partner from the previous breeding attempt and tried to mate with the nest owner. The nest take-over was again unsuccessful and the parasitic female was forced to breed elsewhere with an unpaired male.

Two females laid parasitic eggs during the incubation period of the host. In one case (Case 6, see Results), the female might have made the “best of a bad job”, by laying an egg sired by her social mate in another female’s nest following disastrous circumstances (e.g., after destruction or take-over of her own nest, or after the death of her social mate). In the other case (Case 5, see Results), the parasitic event might have been triggered by the capture of the female on her nest. Note that in Case 5, no other nests in the laying stage were available in the same colony at that moment.

In only one case (Case 2, see Results), the parasitic female may have used a mixed reproductive tactic, laying one parasitic egg on the same day the host female laid her last egg and then continuing to lay in her own nest, which was located in a different barn 40 m away from the parasitized nest. However, we cannot exclude that parasitizing resulted from the female being disturbed, e.g. by farmers or other birds at her own nest.

In eleven nests two eggs appeared within 24 hours interval. From those clutches that could be completely (N = 7) or partially (N = 3) genotyped, only one case of CBP could be confirmed (Case 2 above). Thus, studies reporting CBP in altricial birds that only rely on observations of two eggs appearing within 24 hours (Lyon and Eadie 2017) should be interpreted with caution. Potential counting errors, other mistakes, early nest checks, or females laying later in the day may explain at least some of those cases. For example, in Case 2 described above, the parasitic egg must have been laid after 11:30 (barn swallow females typically lay much earlier in the morning).

False identification of CBP can also arise from otherwise well-executed genetic analysis. In our study, we encountered 14 such cases that did not pass the full CBP verification process. Most of these errors were due to mislabelling of samples in the field or misreading and bad handwriting, leading to the initial misclassification (e.g. Bateman et al. 2013, Marshall et al. 2018). Additionally, we excluded one potential CBP case where a cross-fostering experiment might have caused a misinterpretation of the nestling’s CBP status.

### CBP in other barn swallow studies

The extremely rare occurrence of CBP in our study corresponds to findings from 15 other studies that have analysed parentage in barn swallows using molecular methods and either found no or very few cases of CBP (Table 2). Hasegawa et al. (2010) identified three parasitic nestlings in two nests out of 65 broods (3.1% of broods with CBP) in a population of the Japanese subspecies *Hirundo r. gutturalis*, but they did not identify the parasitic mothers or provide further details. Møller et al. (2003) reported quasi-parasitism (QP), where nestlings were sired by the social male of the focal nest but their mothers were parasitic females, in 5 of 170 broods (2.9%) from a population in Spain. However, it seems that in these cases the entire brood consisted of QP nestlings, suggesting a misidentification of the social mother, as no further details on this extremely rare phenomenon (Griffith et al. 2004) were provided.

**Table 2:** Overview of studies investigating parentage in barn swallow populations using molecular methods. When multiple studies used the same data, only the study with the maximum number of broods and nestlings is presented. In studies where cases of CBP were “not reported”, all offspring samples were classified as either within-pair or extra-pair. Møller a Tegelström (1997) used multi-locus DNA fingerprinting; all other studies performed parentage analysis based on 3-9 microsatellite markers.

| Study | Supspecies | Country | Bredeing seasons(s) | CBP presence | Number of broods | Parasitized broods | Number of nestlings | Parasitic nestlings |
| --- | --- | --- | --- | --- | --- | --- | --- | --- |
| Hasegawa et al. 2010 | H.r. gutturalis | Japan | 2005-2006 | YES - CBP | 65 | 2 | 296 | 3 |
| Kojima et al. 2009 | H.r. gutturalis | Japan | 2006-2007 | NO | 71 | 0 | 301 | 0 |
| Primmer et al. 1995 | H.r. rustica | Denmark | 1988 | NO | 9 | 0 | 44 | 0 |
| Møller a Tegelström 1997 | H.r. rustica | Denmark | 1988-1989 | not reported | 63 | NA | 261 | NA |
| Saino et al. 1997 | H.r. rustica | Italy | 1994 | not reported | 118 | NA | 475 | NA |
| Costanzo et al. 2017 | H.r. rustica | Italy | 2013-2015 | NO | 201 | 0 | 765 | 0 |
| Møller et al. 2003 | H.r. rustica | Spain | 1994 | Yes - QP | 170 | 5 | 674 | 18 |
| Vortman et al. 2013 | H.r. transitiva | Israel | 2009-2010 | not reported | 78 | NA | 328 | NA |
| Kleven et al. 2006 | H.r. erythrogaster | Canada | 2003 | not reported | 86 | NA | 391 | NA |
| Lifjeld et al. 2011 | H.r. erythrogaster | Canada | 2003-2004 | not reported | 139 | NA | 603 | NA |
| Lifjeld et al. 2022 | H.r. erythrogaster | Canada | 2005-2006 | not reported | 147 | NA | 665 | NA |
| Neuman et al. 2007 | H.r. erythrogaster | USA | 2002 | not reported | 53 | NA | 265 | NA |
| Eikenaar et al. 2011 | H.r. erythrogaster | USA | 2009-2010 | not reported | 94 | NA | 424 | NA |
| Hubbard et al. 2015 | H.r. erythrogaster | USA | 2008-2009 | not reported | 157 | NA | 512 | NA |
| Safran et al. 2016 | H.r. erythrogaster | USA | 2009 | not reported | 92 | NA | 411 | NA |
| This study | H.r. rustica | Czech Republic | 2010-2021 | Yes – CBP (QP) | 1945 | 6 (2) | 7816 | 7 (2) |

Based on a study of a Danish population of barn swallows, Møller (1987) suggested that females used CBP as an alternative reproductive strategy. The study concluded that 16.5% of broods (N = 261) contained at least one parasitic egg, based on observations of two eggs being laid in the same nest on one day. The study further suggested that most parasitic eggs were laid by neighbouring females, whereby assignments were based on comparisons of the “odd” egg with the eggs of nearby clutches. However, this subjective method was later challenged by Brown and Sherman (1989), and recent validation demonstrated that human visual identification of parasitic eggs in barn swallow clutches lacks reliability (Hughes et al. preprint). Møller (1987) also attributed parasitic eggs to females whose own clutches were already completed, implying that these females laid parasitic eggs during ongoing incubation. Despite many follow-up studies on the same species (Table 2), no evidence exists for this assertion and we consider it highly unlikely. We are not aware of a single study on any bird species that shows that incubating females continue to lay eggs in the nest of another female. Notably, no further cases of CBP were reported from the Danish population over the following 15 years despite continuous monitoring (Møller and Szép 2005), and genetic studies from later years also failed to detect parasitic nestlings (Primmer et al. 1995, Møller and Tegelström 1997).

### CBP in other Hirundinidae species

CBP has been reported in seven other species of Hirundinidae. Tree swallows (*Tachycineta bicolor*) would seem a prime candidate for higher levels of CBP due to their nesting in secondary cavities, which leads to intense competition for breeding sites and a significant population of female floaters with limited breeding opportunities (Stutchbury and Robertson 1985). For floater females, adopting a CBP strategy could be a last resort, as has been shown or suggested in species like European starling (*Sturnus vulgaris*; Sandell and Diemer 1999) and spotless starling (*Sturnus unicolor*; Redondo et al. 2022). However, contrary to these expectations, CBP in tree swallows has only been reported in early studies based on the less reliable criterion of “two eggs in the nest within 24 hours” (Kuerzi 1941, Lombardo 1988). Cases of CBP were extremely rare in studies that used parentage analysis (Barber et al. 1996, Kempenaers et al. 1999, Conrad et al. 2001, Bitton et al. 2007).

Genetic studies have suggested that CBP may serve as an alternative reproductive strategy in another cavity-nesting species, the purple martin (*Progne subis*). Morton et al. (1990) found that 36% of nestlings (N = 28) cared for by second-year females were parasitic, while Tarof et al. (2011) identified 4.7% of all nestlings (N = 1235) across almost 300 broods as parasitic. Neither of these studies was able to identify the parasitic females. A study on a population of sand martins (*Riparia riparia*) suggested seven cases of quasi-parasitism (N = 167 nestlings), but the authors admitted the possibility of misassignment (Alves and Bryant 1998). In another study on this species, Hoogland and Sherman (1976) reported 9 putative CBP cases identified on the basis of the “two eggs laid in 24 hours” criterion, but they clearly stated that these cases were likely caused by "delays in the timing of egg deposition or to variations in the timing of nest examination”.

Several studies investigated CBP in four species of cliff-nesting swallows from the genus *Petrochelidon*, some using the “two eggs per day” criterion. However, these species build inaccessible domed nests e.g. on cliffs, under bridges, or in culverts (Brown and Brown 1998). Accessing these nests for direct checks or nestling sampling is challenging, often requiring specialized equipment or even the construction of artificial side entrances (Earlé 1986, Magrath et al. 2002, Weaver and Brown 2004). The only two genetic studies conducted on the fairy martin (*Petrochelidon ariel*), reported only a single case of CBP based on protein analysis of 60 eggs (Manwell and Baker 1975), or no CPB based on 465 nestlings from 169 nests (Magrath et al. 2002). Hammers et al. (2009) also found no evidence of CBP in fairy martins based on the “two egg per day” criterion. CPB was suspected in the South African cliff swallow (*Petrochelidon spilodera*) based on unusually large clutches of four eggs (Earlé 1986) and in the cave swallow (*Petrochelidon fulva*), based on 60 clutches with two eggs laid within 24 hours (4.8%, N = 1252; Weaver and Brown 2004). In the cliff swallow, Brown and Brown (1989) witnessed CBP at least 30 times, with parasitic females entering host nests and laying eggs rapidly, similar to the behaviour of obligate brood parasites (Gloag et al. 2013, Jelínek et al. 2021). Often, these parasitic females had their own nest and were direct neighbours (Brown 1984). They also observed two instances in which a female physically transported an egg to another cliff swallow nest (Brown and Brown 1988).

## Conclusions

In sum, although cases of CBP have been reported in several Hirundinidae species, compelling evidence supporting the hypothesis that it is used as an alternative reproductive strategy is missing, with the exception of the cliff swallow. In this species, CBP is well-documented, being observed at high rates (9.9% of nests, N = 4942, Brown and Brown 1989; 11.4%, N = 351, Weaver and Brown 2004), and with numerous direct observations of parasitic behaviour. In other species, relatively high CBP rates have been reported based on traditional criteria (e.g., two eggs appearing in a nest within 24 hours), but these findings were either not confirmed by genetic studies (e.g. barn swallow and tree swallow) or no genetic studies have been conducted (e.g. South African cliff swallow and cave swallow). Parentage analyses thus suggest that the “two eggs laid within 24 hours” criterion is unreliable (see also our results). To increase our understanding of CBP, future research should focus on identifying both parents of the parasitic eggs. Only when the identity of both parents is known, we can fully unravel the circumstances that drive females to dump an egg in the nest of a conspecific. To our knowledge, our study is the first to achieve this in a species of the Hirundinidae. We demonstrated that the circumstances surrounding CBP in our barn swallow population deviated from the standard breeding situations, and included (failed) nest take-overs, human disturbance, and the disappearance of a social partner as potential triggers for CBP.

## Funding

This study was financially supported by a project from the Czech Science Foundation (grant numbers 20-06110Y, 21-22160S and 25-17505S) and by the Max Planck Society (to SK).

## Conflict of interest

Authors declare no conflict of interests.

## Ethical approval

We declare that the sampling was approved by the animal and ethics representatives of The Czech Academy of Sciences and nature conservation authorities (62065/2017-MZE-17214 and MZP/2020/630/964).

## ACKNOWLEDGEMENTS

We thank all of our field assistants especially František Buben, Kristýna Míčková, Lukáš Pazdera and Martina Němcová, née Soudková and lab technicians Radka Valterová, Alena Fornůsková, and Kamila Vlčková. We also thank Bart Kempenaers for valuable comments and carefully editing the manuscript. This study would not have been possible without the intense collaboration of local farm owners, specifically the Kotrba family at Hamr farm, the Kraus and Pulec families at the Šaloun site, Jan Kačerovský and the staff of the Břilice and Stará Hlína farms and all house owners from the Chotovice village. We also thank local conservation authorities for permission to conduct the fieldwork.

## Notes

### Competing Interest Statement

The authors have declared no competing interest.

### Summary of Updates

We improved wording for the clarity of the text.

