## Supplementary material for "Conspecific brood parasitism in the barn swallow (*Hirundo rustica*) and other Hirundinidae": Table S1 (from another version of this preprint: version 2, 2025-02-21)

**Table S1:** Characteristics of 17 microsatellite markers used in this study. Given are the locus name, the primer sequence and dye, the multiplex mix to which it belonged, concentration of each primer in micro molar (C), the number of alleles (k, based on information from all genotyped individuals). Following characteristics are depicted as minimum and maximum values from all 15 parentage analyses performed to assign maternity: - expected and observed heterozygosity (H_e_ and H_o_), the estimated frequency of null alleles (F_null_), the non-exclusion probability for the first parent (NE-1P), and the non-exclusion probability for the second parent (NE-2P).

| **Loci** | **Primer sequences (5’ - 3’) and dye** | **Mix** | **C (μM)** | **k** | **marker range (bp)** | **H_e_** | **H_o_** | **F_null_** | **NE-1P** | **NE-2P** | **Reference** |
| --- | --- | --- | --- | --- | --- | --- | --- | --- | --- | --- | --- |
| HrU5 | F: 6FAM-TCAACAAGTGTCATTAGGTTC | 1 | 0.34 | 41 | 110–144 | 0.904 – 0.916 | 0.886 – 0.915 | -0.0032 – 0.0145 | 0.289 – 0.323 | 0.169 – 0.193 | Primmer et al. 1995 |
|  | R: AACTTAGATAAGGAAGGTATAT |  |  |  |  |  |  |  |  |  |  |
| TG11-011 | F: 6FAM-ACAAACTAAGTACATCTATATCTGAAG | 1 | 1.28 | 14 | 209–233 | 0.768 – 0.81 | 0.750 – 0.822 | -0.0074 – 0.0232 | 0.547 – 0.616 | 0.371 – 0.434 | Dawson et al. 2010 |
|  | R: TAAATACAGGCAACATTGG |  |  |  |  |  |  |  |  |  |  |
| 2F9 | F: VIC-GCATTTCTGGGCTGTAACAT | 1 | 0.84 | 22 | 78–110 | 0.892 – 0.913 | 0.888 – 0.922 | -0.0186 – 0.0061 | 0.302 – 0.364 | 0.177 – 0.222 | Heber et al. 2013 |
|  | R: AAAGGACAATGTAATTGGTG |  |  |  |  |  |  |  |  |  |  |
| Hir10 | F: VIC-GGACAAGGGGAGTCTT | 1 | 0.18 | 15 | 150–202 | 0.827 – 0.849 | 0.773 – 0.852 | -0.0095 – 0.0357 | 0.471 – 0.518 | 0.305 – 0.345 | Tsyusko et al. 2007 |
|  | R: ATTCAGCCAGCCTCTAAT |  |  |  |  |  |  |  |  |  |  |
| Hir11 | F: NED-AACACCTGAAAACCTACAC | 1 | 0.16 | 29 | 164–247 | 0.808 – 0.839 | 0.782 – 0.867 | -0.0207 – 0.0229 | 0.492 – 0.550 | 0.323 – 0.374 | Tsyusko et al. 2007 |
|  | R: CTTTGAGCAAAATGAGTG |  |  |  |  |  |  |  |  |  |  |
| Tgu07 | F: PET-CTTCCTGCTATAAGGCACAGG | 1 | 0.40 | 17 | 90–124 | 0.792 – 0.812 | 0.780 – 0.852 | -0.0277 – 0.0075 | 0.546 – 0.580 | 0.370 – 0.403 | Slate et al. 2007 |
|  | R: AAGTGATCACATTTATTTGAATAT |  |  |  |  |  |  |  |  |  |  |
| TG11-000 | F: PET-TTGCTACCARAATGGAATGT | 1 | 1.20 | 17 | 233–265 | 0.789 – 0.809 | 0.771 – 0.825 | -0.0173 – 0.0187 | 0.534 – 0.568 | 0.358 – 0.390 | Dawson et al. 2010 |
|  | R: TCCTAACCATGAGAAGCAGA |  |  |  |  |  |  |  |  |  |  |
| 3007/3112 | F: PET-TACATACAGGCTCTACTCCT | 1 | 1.00 | 4 | 342–354 | 0.380 – 0.443 | 0.488 – 0.558 | -0.1424 – -0.1132 | 0.901 – 0.929 | 0.789 – 0.838 | Ellegren and Fridolfsson 1997 |
|  | R: CCCCTTCAGGTTCTTTAAAA |  |  |  |  |  |  |  |  |  |  |
| Gf16 | F: 6FAM-CCCTTCAGGGCATGAGTGAGG | 2 | 0.18 | 14 | 103–130 | 0.690 – 0.731 | 0.620 – 0.739 | -0.0065 – 0.0580 | 0.686 – 0.738 | 0.514 – 0.574 | Petren 1998 |
|  | R: ATGTCATGAACTCAACCAACTCC |  |  |  |  |  |  |  |  |  |  |
| ADCYAP1_bm | F: 6FAM-GATGTGAGTAACCAGCCACT | 2 | 0.32 | 25 | 145–184 | 0.788 – 0.853 | 0.769 – 0.876 | -0.0143 – 0.0192 | 0.453 – 0.576 | 0.291 – 0.399 | Steinmeyer et al. 2010 |
|  | R: ATAACACAGGAGCGGTGA |  |  |  |  |  |  |  |  |  |  |
| IND40 | F: 6FAM-ACCGAAACAACAGAAACAGT | 2 | 0.60 | 26 | 222–252 | 0.745 – 0.824 | 0.726 – 0.822 | -0.0173 – 0.0182 | 0.495 – 0.624 | 0.325 – 0.440 | Sefc et al. 2001 |
|  | R: AGAACGCTAAGTGAATGTCC |  |  |  |  |  |  |  |  |  |  |
| Z37B | F: VIC-AACTGGTTGTAGGTATAGTGCAATTATG | 2 | 0.28 | 15 | 85–135 | 0.770 – 0.794 | 0.843 – 0.883 | -0.0674 – -0.0347 | 0.578 – 0.615 | 0.399 – 0.435 | Dawson et al. 2015 |
|  | R: GATTACAAAGCCAATATGGATGC |  |  |  |  |  |  |  |  |  |  |
| PiJ14-23-CEST | F: VIC-ATCTGGCATKGAAAACTTGG | 2 | 0.26 | 18 | 156–190 | 0.846 – 0.861 | 0.823 – 0.887 | -0.0239 – 0.0178 | 0.441 – 0.474 | 0.280 – 0.307 | Olano et al. 2010 |
|  | R: CTCCCTGCACCCCAAAC |  |  |  |  |  |  |  |  |  |  |
| Hir20 | F: VIC-GAAGTTGGAGAAAGATTAG | 2 | 0.26 | 63 | 220–333 | 0.819 – 0.880 | 0.768 – 0.869 | -0.0047 – 0.0343 | 0.382 – 0.527 | 0.236 – 0.355 | Tsyusko et al. 2007 |
|  | R: TTATTGCTCTGGGTATGT |  |  |  |  |  |  |  |  |  |  |
| LEI160 | F: NED-GCAGACAGCCGTTAATATATGCG | 2 | 0.32 | 16 | 158-188 | 0.827 – 0.859 | 0.792 – 0.868 | -0.0096 – 0.0208 | 0.443 – 0.497 | 0.283 – 0.326 | Gibbs et al. 1997 |
|  | R: AACCAAAACACAAGCTCTTGCA |  |  |  |  |  |  |  |  |  |  |
| Hir22 | F: NED-ATCCGCACCTAATGT | 2 | 0.24 | 52 | 234–313 | 0.899 – 0.906 | 0.842 – 0.911 | -0.0077 – 0.0364 | 0.312 – 0.331 | 0.186 – 0.199 | Tsyusko et al. 2007 |
|  | R: GATTCACATATCCCATCTAG |  |  |  |  |  |  |  |  |  |  |
| Hir19 | F: PET-GCTCACAACCAGCTAGAC | 2 | 0.16 | 31 | 130–214 | 0.849 – 0.888 | 0.738 – 0.861 | 0.0081 – 0.0714 | 0.367 – 0.461 | 0.225 – 0.298 | Tsyusko et al. 2007 |
|  | R: ATAGCCACAGGGAAAGTCT |  |  |  |  |  |  |  |  |  |  |

Dawson DA, Horsburgh GJ, Küpper C, Stewart IRK, Ball AD, Durrant KL, Hansson B, Bacon I, Bird S, Klein A, Krupa AP, Lee JW, Martín-Gálvez D, Simeoni M, Smith G, Spurgin LG, Burke T (2010) New methods to identify conserved microsatellite loci and develop primer sets of high utility – as demonstrated for birds. Mol Ecol Resour 10:475–494

Dawson DA, Brekke P, Dos Remedios N, Horsburgh GJ (2015) A marker suitable for sex-typing birds from degraded samples. Conserv Genet Resour 7:337–343

Ellegren H, Fridolfsson AK (1997) Male-driven evolution of DNA sequences in birds. Nat Genet 17:182–184

Gibbs M, Dawson DA, McCamley C, Wardle AF, Armour JAL, Burke T (1997) Chicken microsatellite markers isolated from libraries enriched for simple tandem repeats. Anim Genet 28:401–417

Heber S, Varsani A, Kuhn S, Girg A, Kempenaers B, Briskie J (2013). The genetic rescue of two bottlenecked South Island robin populations using translocations of inbred donors. Proc R Soc B 280: 1752

Olano-Marin J, Dawson DA, Girg A, Hansson B, Ljungqvist M, Kempenaers B, Mueller JC (2010) A genome-wide set of 106 microsatellite markers for the blue tit (*Cyanistes caeruleus*). Mol Ecol Resour 10:516–32

Petren K (1998) Microsatellite primers from *Geospiza fortis* and cross-species amplification in Darwin's finches. Mol Ecol 7:1782–1784

Primmer CR, Møller AP, Ellegren H (1995) Resolving genetic relationships with microsatellite markers: a parentage testing system for the swallow *Hirundo rustica*. Mol Ecol 4:493–498

Sefc KM, Payne RB, Sorenson MD (2001) Characterization of microsatellite loci in village indigobirds *Vidua chalybeata* and cross-species amplification in estrildid and ploceid finches. Mol Ecol Notes 1:252–254

Slate J, Hale MC, Birkhead TR (2007) Simple sequence repeats in zebra finch (*Taeniopygia guttata*) expressed sequence tags: a new resource for evolutionary genetic studies of passerines. BMC Genomics 8:52

Steinmeyer C, Mueller JC, Kempenaers B (2010) Search for informative polymorphisms in candidate genes: clock genes and circadian behaviour in blue tits. Genetica 136:109–117

Tsyusko OV, Peters MB, Hagen C, Tuberville TD, Mousseau TA, Møller AP, Glenn TC (2007) Microsatellite markers isolated from barn swallows (*Hirundo rustica*). Mol Ecol Notes 7:833–835
